# Anatomy-aware, label-informed approach improves image registration for challenging datasets

**DOI:** 10.1101/2025.08.11.669599

**Authors:** Rachel A. Roston, Nicholas J. Tustison, A. Murat Maga

**Affiliations:** Center for Developmental Biology and Regenerative Medicine, Seattle Children’s Research Institute, Seattle, Washington, USA; Department of Radiology and Medical Imaging, University of Virginia, Charlottesville, Virginia; Department of Pediatrics, University of Washington, Seattle, Washington, USA

## Abstract

Image registration-based volumetric morphometrics have emerged as a valuable method for identifying subtle morphological differences in neuroimaging and other biomedical images. However, accurate registration out-of-the-box remains challenging when overt morphological phenotypes—such as those observed in developmental and comparative studies—are present in a dataset. A new label-informed image registration function developed in the ANTsX ecosystem provides an easy to use, generalizable solution for anatomy-aware registration of a wide diversity of morphological variation. In this approach, segmentations (*i.e.*, labels) provide *a priori* regional correspondences that guide the registration. These labels can be generated by any method–manually, using semi-automated tools, or through deep learning-based approaches–and allow morphological experts to define regions of correspondence based on biological concepts of homology (*e.g.,* tissue origin, gene expression patterns). Here we demonstrate the utility of this label-informed image registration approach for improving the registration knockout mouse embryos which fail to register to a wildtype (normative) template image by traditional registration methods. E15.5 *Gli2^−/-^* mouse embryos show a severe scoliosis and radical topological rearrangement of the internal organs. Compared to traditional, intensity-only registration, the new label-informed image registration improved the correspondence of knockout subjects to the canonical template image, which resulted in increased power and sensitivity of downstream statistical analyses. All in all, label-informed image registration provides a flexible and customizable method to allow image registration in datasets for which registration-based morphometrics were previously unfeasible, unlocking new potential applications of registration-based morphometrics in developmental, comparative, and evolutionary studies.

## Introduction

Morphometrics–the mathematical and statistical study of form–is an essential part of studying biological variation (Bateson 1894; Thompson 1917). Uncovering the origins and causes of morphological variation requires the ability to quantify phenotypic differences despite overt differences in morphology. Quantitative morphometrics provide important insights into the developmental, physiological, and evolutionary mechanisms that produce qualitative phenotypes (Cooper & Albertson 2008). Many, if not most, qualitative phenotypic differences begin as quantitative differences at earlier developmental timepoints or lower hierarchical levels of biological organization (*e.g.*, cell signaling, molecular interactions) (Hallgrímsson et al. 2012).

Over the past several decades, comparative morphometrics methods have shifted from traditional two-dimensional measurements with calipers to incorporate a range of 2D and 3D geometric methods inspired by D’Arcy Thompson’s (1917) deformation diagrams (Dryden & Mardia 2016; Goswami & Clavel 2025; Roth 1993; Zelditch et al. 2012). Advances in imaging and surface scanning, software, and computational methods have made it easier than ever to collect, analyze, and interpret morphometric data for a wide array of biological questions (*e.g.*, Avants et al. 2011; Boyer et al. 2015; Porto et al. 2021; Rolfe et al. 2021; Tustison et al. 2021; Zhang et al. 2022). Numerous tools and methods have been developed to analyze diverse morphologies by comparing landmark point configurations and surfaces, but volumetric (or, voxel-based) morphometric approaches have notably been missing in many studies of biological variation.

In voxel-based morphometrics, image registration maps each voxel in a 3D volumetric image (*e.g.,* CT scan, MRI) to a template image, providing dense volumetric shape information that cannot be captured with landmark- or surface-based approaches (Chen et al. 2023; Deserno 2001). Intensity image registration calculates the transformations required to align the subject image with the template (Chen et al. 2023; Deserno 2001). Then, statistical tests are performed on the transformed images or on the deformation fields (*i.e.,* tensors, transforms) to compare subjects and identify statistically significant quantitative differences in morphology (*e.g.*, (Ashburner & Friston 2000; Horner et al. 2021; Wong et al. 2014). The output of these statistical tests may be visualized as heatmaps or plots of principal components (*e.g.*, Ashburner & Friston 2000; Horner et al. 2021; Wong et al. 2014).

While the methods for intensity image registration, voxel-based morphometry (VBM), and tensor-based morphometry (TBM) have existed for over two decades, their application has been primarily restricted to quantifying subtle morphological differences in neuroimaging. In these applications, morphological variation is often limited by the design of a study. Studies using VBM and TBM have focused primarily on inter-subject differences in a single species and developmental stage, multi-modal imaging of a single individual, such as aligning CT and PET scans for the same patient, or longitudinal studies following a single individual. Such studies have provided valuable insights on a vast array of biological questions and pathologies.

VBM and TBM have been proposed as methods for high-throughput, automated phenotype detection and quantitative analysis in high-throughput genetic screens, but have not been widely integrated as standard approaches in the characterization of mutant mouse phenotypes (Dickinson et al. 2016; Horner et al. 2021; Wong et al. 2014; Zamyadi et al. 2010). An ongoing challenge is that existing image registration algorithms presume a certain amount of similarity in the images being registered. However, many mutations cause overt or gross phenotype differences such as large differences in the size, shape, orientation, or position of anatomical structures, and structures that are missing entirely (Dickinson et al. 2016; Groza et al. 2023; Horner et al. 2021).

Overt phenotypes remain difficult to register with current methods for several reasons. Registration generally proceeds in a hierarchical manner, starting with gross alignment of the images through linear transformations (*i.e.,* translation, rotation, scaling, and shear) and then fine-tuning image alignment with local transformations (Chen et al. 2023; Deserno 2001; Horner et al. 2021). Local transformations are constrained by registration parameters to minimize the distance of transformations, reduce registration times, and decrease memory requirements. This approach presumes that the initial linear transformations produce a relatively good initial alignment of the images, and fails if subjects have different overall shapes that cannot be approximately aligned through linear transformations (*e.g.*, differences in posture). Testing different registration parameters to optimize an automated registration algorithm is time- and memory-intensive. And, incomplete penetrance, variable expression, and small sample sizes in reverse genetic screens mean that optimal parameters for one subject may not apply to another subject with the same genetic mutation (Horner et al. 2021). Also, the whole-body surveys preferred for phenotype discovery require simultaneous registration of multiple anatomical structures, all of which can vary in size, shape, position, and orientation, and in their topological inter-relationships. Registering multiple anatomical structures within an image, such as multiple organs in the whole body, therefore has more degrees of freedom than registering a single organ, such as the brain. And, as a result, selecting registration parameters requires balancing good alignment of certain structures with misalignment of other structures (Horner et al. 2021).

Here we present an anatomy-aware, label-informed image image registration approach available in the open-access, open-source ANTsX ecosystem that allows the registration of a broader range of morphological phenotypes, and therefore VBM and TBM studies of subjects with overt phenotypes (Avants et al. 2009; Avants et al. 2011; Tustison et al. 2021). Traditional medical image registration approaches find anatomical correspondences based on intensity values and geometry (Chen et al. 2023; Deserno 2001). This approach works well when only a single organ or tissue is in the field of view and when the geometry is relatively similar between subjects. In whole body scans with multiple organs and tissues, both the intensity values and internal geometry can vary substantially. Segmentation is the process of labeling each voxel in an image to identify it as part of a larger region of interest. labels (*i.e.*, labels) may comprise a region or lobe within an organ, the whole organ, spaces outside organs, or a larger region of interest such as the thoracic or abdominal cavity.

Using labels to initialize intensity image registration allows experts in morphology to guide and customize image registration according to the specific goals and questions of a study. Corresponding regions may be identified based on expert knowledge of developmental origins, gene expression, or other criteria relevant to a study; this information may not be readily apparent or contained within an image itself. Morphologists are therefore able to choose which features or regions should be considered homologous and compared based on biological concepts of homology (*e.g.*, Abouheif 1997; Patterson 1982; Roth 1984; Roth 1991; Roth 1993; Rutishauser & Moline 2005; Scotland 2010; Shubin et al. 2009; Van Valen 1982; Wagner 2014). Labels can also allow customization of which regions of interest should be prioritized in the registration. For example, a missing structure (*e.g*., a missing heart) can cause neighboring structures (*e.g*., lungs) to radically shift in position. While the missing structure itself cannot be registered, label-informed registration can support the registration of the neighboring structures despite their radical shift in position.

To demonstrate how label-informed image registration using the labelImageRegistration function in ANTsR affects registration results and downstream statistical analyses, we have selected a genetic knock-out line as a test case that has failed to register to a wildtype-based template image using existing intensity-only registration methods. GLI2 is a transcription factor involved in transduction in the sonic hedgehog (SHH) signaling pathway (Park et al. 2000). Knocking out *Gli2* causes multiple overt phenotypes across the whole embryo, including developmental anomalies in the brain, face, spinal cord, vertebral column, heart, lungs, digestive system, and urogenital system (Park et al. 2000) (Figure 1). Visual observation suggests that these embryos also have several quantitative phenotypic differences, such as a reduction in bladder volume and alterations in liver shape. Measuring these quantitative differences requires accurate image registration despite the overt phenotypic differences that affect overall embryo shape and topological differences.

**Figure 1.**
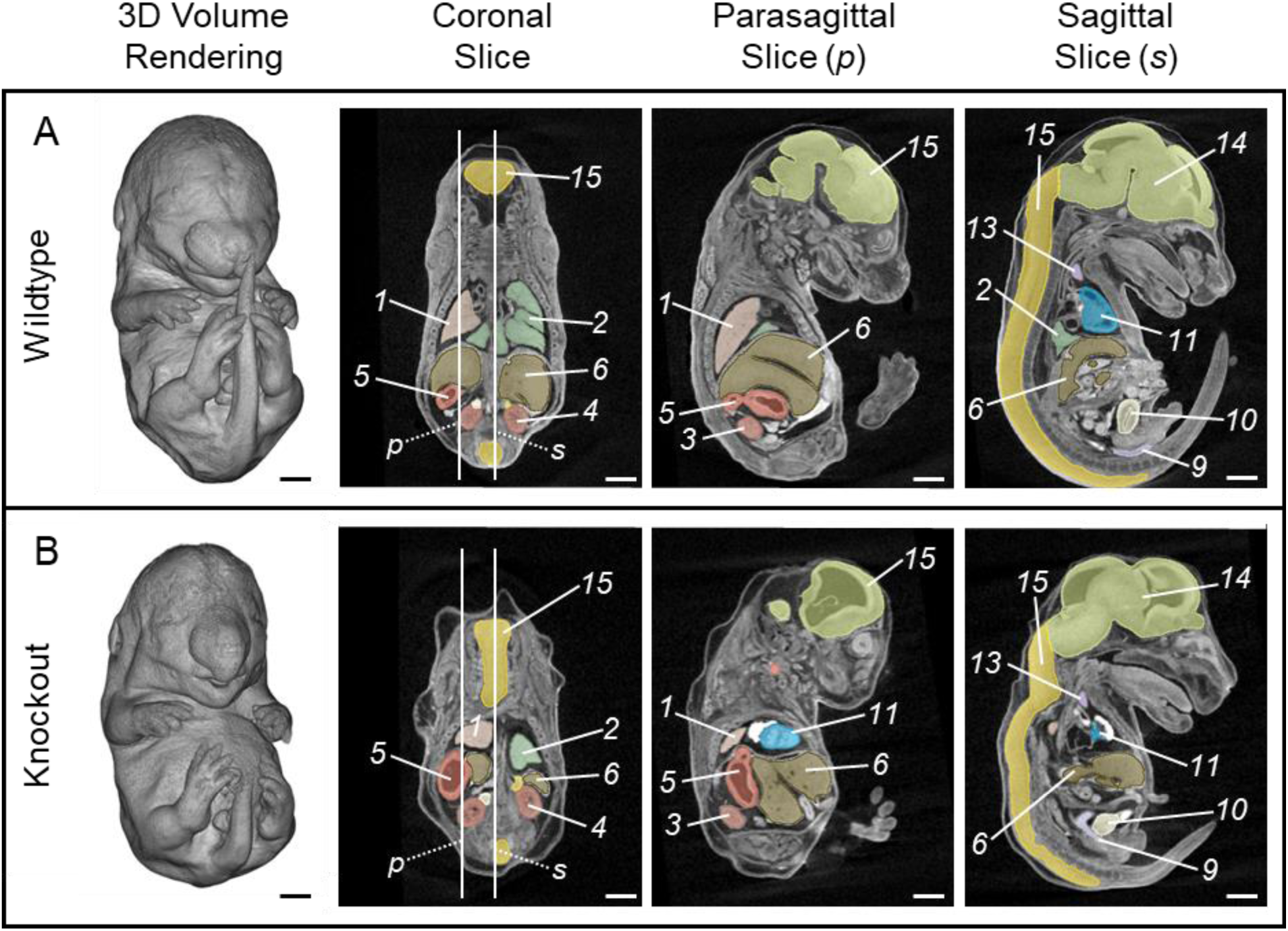
Representative diceCT scans showing organ topology in (A) a representative wildtype (Gli2^+/+^; Scan_0093) and (B) a representative knockout (Gli2^−/-^; Scan_0105) subject. Vertical lines in the coronal slice show the locations of the parasagittal (*p*) and sagittal (*s*) slices for each subject. Scale bar = 1 cm. Numbers indicate organ label: *1 left lung, 2 right lung, 3 left kidney, 4 right kidney, 5 stomach, 6 liver, 7 left adrenal gland, 8 right adrenal gland, 9 rectum, 10 bladder, 11 heart ventricles, 12 left thymic rudiment, 13 right thymic rudiment, 14 brain, 15 spinal cord*.

Our results show that label-informed image registration produces qualitatively better alignment of *Gli2* knockout embryos over the wildtype template image than intensity-only registration. And, this improved alignment increases the power of downstream statistical analyses and their interpretability. These results suggest that a label-informed approach to image registration is a broadly applicable and customizable method that allows volumetric (voxel- and tensor-based) morphometrics for a wider range of phenotypic variation. Thus, label-informed image registration can allow broader application of volumetric morphometric approaches in developmental and comparative contexts where it previously was minimally informative.

## Methods

### Embryo Collection and Micro-CT

Nine *Gli2^+/+^*and nine *Gli2^−/-^* mouse embryos were collected on embryonic day 15.5 (E15.5) by S. Gombart and E. Vincent at Seattle Children’s Research Institute in accordance with IACUC protocol 00030. These subjects were a subset of over 250 embryos collected and scanned for an ongoing reverse genetic screen. *Gli2* was selected for the present study because *Gli2^−/-^* embryos have a severe, overt phenotype that failed to register accurately to a wildtype template image using existing intensity image registration methods (Figures 1-3).

Subjects were fixed overnight in 4% paraformaldehyde solution (PFA) and stored in PBS at 4°C until preparation for scanning. To stabilize the soft tissues, subjects were embedded in hydrogel following Wong et al. (2013) and Dickinson et al. (2016) (Dickinson et al. 2016; Wong et al. 2013). To summarize, hydrogel solution containing 4°C 4% PFA (16% PFA in aqueous solution, VWR), 4°C 4% acrylamide (Fisher Scientific), 4°C 0.05% bis-acrylamide (Fisher Scientific), 0.25% VA-044 (FUJIFILM Wako Chemicals), 0.05% saponin (Sigma Life Science), and 4°C PBS was mixed on ice and frozen in ∼12-mL aliquots. In preparation for scanning, fixed specimens were transferred from PBS to liquid (thawed) hydrogel solution. After 70-74 hours of incubation at 4°C, hydrogel was polymerized at 37°C for three hours and excess hydrogel was carefully removed from the outside of the specimens. The specimens were stained in 0.1 N iodine solution (Sigma-Aldrich) for 24 hours. After 24 hours, specimens were washed in room temperature PBS twice to remove excess iodine from the outside of the subjects and embedded in 1% agarose to stabilize them for microCT scanning. 3D scanning of subjects was conducted at the SCRI MicroCT Imaging Facility (<u>RRID:SCR_024678</u>), using Bruker Skyscan 1272 microCT that was funded by an NIH shared instrument grant (<u>S10OD032302</u>). Specimens were scanned with isotropic spacing of 9um^3^ (frame averaging = 5) or 18um^3^ (frame averaging = 3), a 1-mm aluminum filter, and vertical rotation of 180 degrees. DiceCT images acquired at 9um were resampled to 18um^3^ spacing prior to further analysis using the batch mode in the SkyscanRecontImport module from SlicerMorph (Rolfe et al. 2021).

### Template Image

The template image for this study was generated by averaging thirty-five wildtype micro-CT volumes from eight inbred and outbred knockout mouse lines (Table 1). The most common strain backgrounds were C57BL/6J and C57BL/6NCrl. First, all subjects were rigidly aligned to a common orientation that is congruent with the standard anatomical planes. An initial linear average of the rigidly-aligned subjects was used to initialize the template building process. To build the template, all subjects were aligned with the initial linear average using both linear and deformable registration. Then, a new template image was created by averaging all the subjects in the new space and applying filters (*e.g.*, sharpening and smoothing). The process then repeated iteratively using the template image calculated from the previous iteration. A total of three iterations was used to obtain an anatomically correct template from the selected wildtype subjects. More information about ANTs template building process can be found in the ANTsX Github documentation (Avants et al. 2011, (https://github.com/ANTsX/ANTs/blob/master/Scripts/antsMultivariateTemplateConstruction2.sh).

**Table 1.** Knock-out lines and strain backgrounds used to build the normative (wildtype) template image.

| Knockout Line | Strain Background | Number (n) Used in Building the Template Image |
| --- | --- | --- |
| *Gli1*<sup>tm2Alj</sup>/J | Outbred: 35.3-41.2% 129S | 5 |
| *Gli2*<sup>tm6Alj</sup>/J | Outbred: 89.1-91.7% C57BL/6J | 4 |
| *Gli3*<sup>tm1Alj</sup>/J | Outbred: 89.1-91.7% C57BL/6J | 1 |
| *Sptan1*<sup>em1(IMPC)Mbp</sup>/Mmucd | Close to inbred: C57BL/6NCrl | 9 |
| *Tbx1*<sup>em1(IMPC)Mbp</sup>/Mmucd | Close to inbred: C57BL/6NCrl | 8 |
| *Tcf21*<sup>em1(IMPC)Mbp</sup>/Mmucd | Close to inbred: C57BL/6NCrl | 4 |
| *Ttc21b*<sup>aln</sup> | Close to inbred: B6.A(FVB) | 1 |
| *Vangl2*<sup>em1(IMPC)Mbp</sup>/Mmucd | Close to inbred: C57BL/6NCrl | 3 |

### Segmentation

Fifteen organ labels (*i.e.,* segments) were selected by combining labels used in the published IMPC atlas and MEMOS (Rolfe et al. 2023; Wong et al. 2012). Segments labeled the following organs: brain, spinal cord, thymic rudiments (2), heart ventricles, lungs (2), liver, stomach, kidneys (2), adrenal glands (2), bladder, rectum. Manual segmentation of individual organs and the whole subject mask was conducted using the segmentation tools in 3D Slicer and SlicerMorph (Fedorov et al. 2012; Kikinis et al. 2014; Rolfe et al. 2021).

### Registration

Prior to this study, intensity images were downsampled from 18um^3^ to 60um^3^ spacing to allow data-sharing and to decrease the memory requirements for running the registration scripts provided in the Github repository for this project. Subject images were also rigidly aligned with the template prior to this study to support easy and rapid visualization of subject anatomy in standard anatomical planes.

To compare intensity-only registration and label-informed intensity registration approaches, subjects were registered to the template image using the **antsRegistration()** and **labelImageRegistration()** functions in the ANTsR package (Avants et al. 2011; Avants 2024; Kandel et al. 2024; Tustison et al. 2021). The **labelImageRegistration()** function enables anatomically-aware registration by integrating anatomical label maps and optional intensity image pairs into the optimization process. In this study, both label maps and intensity image pairs were used. The registration proceeds in a flexible multi-stage framework: (1) initialization using either a user-supplied transform or a linear alignment (*e.g*., rigid, affine) based on the centers of mass of corresponding label maps; (2) deformable registration using intensity-based similarity metrics (typically via the SyN algorithm), initialized by the output of the first stage; and (3) optional refinement using a label-based similarity metric. All label images remain fixed during optimization and serve as biologically meaningful anchors that guide deformation, and individual labels may be differentially weighted to emphasize structures of interest. In the intensity-only pipeline, the linear initialization was derived from direct registration of the intensity images; in contrast, the label-informed pipeline initialized alignment using center-of-mass alignment of the organ label maps. Both approaches then proceeded through deformable registration using intensity image similarity. All registrations excluded non-subject voxels using manually segmented whole-subject masks. Importantly, the same functionality is available in the Python-based ANTsPy library via the **label_image_registration()** function. Registration scripts and associated image data are available at https://github.com/raroston/labelImageRegistration_Comparison.

### Statistical Analyses

#### Organ Volume Analysis

Physical volumes of labels were calculated using the **labelStats()** function of ANTsR and t-tests were performed using base R functions package (Avants et al. 2011; Avants 2024; Kandel et al. 2024; R Core Team 2023; Tustison et al. 2021).

#### Tensor Based Morphometry (TBM)

Jacobian determinants were calculated from the forward transforms generated from each registration approach using the **CreateJacobianDeterminantImage()** function of ANTsR (Avants et al. 2011). For both registration approaches, a voxel-wise linear regression was performed to compare the Jacobian determinants from wildtype and knockout subjects. The p-values were adjusted for multiple comparisons using false discovery rate (FDR, a < 0.05) by using the relevant functions in the ANTsR and base R libraries (Avants et al. 2011; Avants 2024; Kandel et al. 2024; R Core Team 2023; Tustison et al. 2021). Heatmaps of significantly different voxels were visualized in 3D Slicer (Fedorov et al. 2012; Kikinis et al. 2014).

#### Principal Component Analysis (PCA)

PCA was conducted on combined inverse transforms using the **multichannelPCA()** function in ANTsR (Avants et al. 2011; Avants 2024). To create visually discernable outputs, transforms associated with each PC axis, eigenvectors were arbitrarily scaled by -500 (negative PC extrema) and 500 (positive PC extrema) and used to transform the template image into subject space.

## Results

### Severe scoliosis in *Gli2^−/-^* subjects

*Gli2^−/-^* (knockout) subjects show severe scoliosis with sharp bends and twists in the spinal cord and misshapen or missing vertebrae, causing the torso to appear longitudinally compressed and relatively wide (Figure 2). Compared to *Gli2^+/+^* (wildtype) subjects, the cranial base of the knockout subjects has a less acute angle with the overall body axis. Because of these differences in embryonic curvature, the relative orientations of large body regions, and the resulting relative displacement of internal organs, the knockout subjects do not register easily to a wildtype template image (Figure 3A).

**Figure 2.**
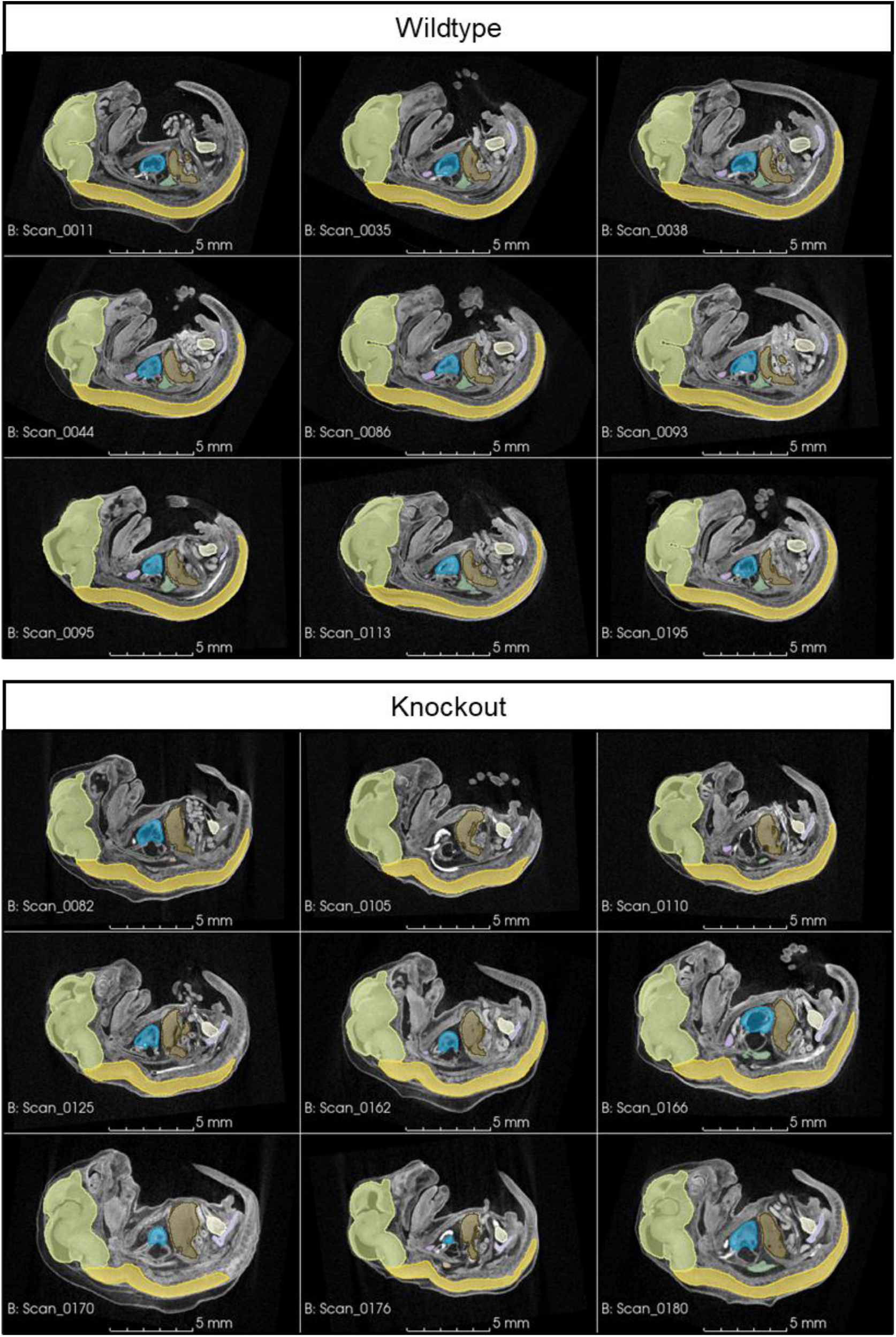
Gross morphological differences in knockout embryos compared to wildtype embryos. The spinal cord and vertebral columns of *Gli2^−/-^* (knockout) embryos have severe bends and twists (*i.e.,* scoliosis), which alters the overall body shape and topology of internal organs.

**Figure 3.**
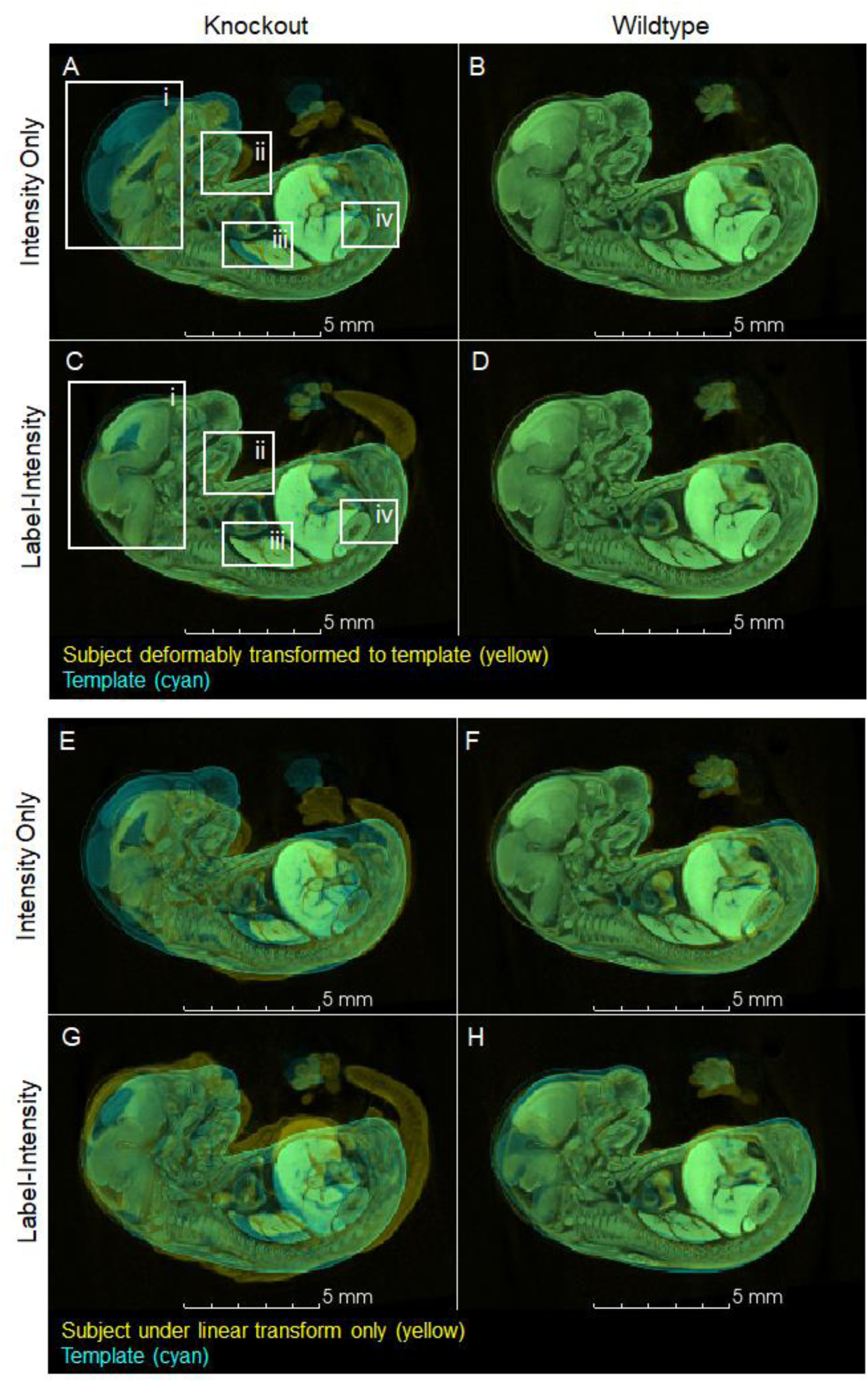
Intensity-only and label-intensity registration results for a representative knockout (*Gli2^−/-^*; Scan_0105) and a representative wildtype subject (*Gli2^+/+^*, Scan_0093) (yellow) overlaid over the template image (cyan). (A-D) Subjects transformed to template space using combined forward transforms. (E-H) Subjects under the initial linear transform only. For the knockout subject, intensity-only registration (A) resulted in a poor overlay of the brain (i), lower jaw (ii), lung (iii), kidney (iv), and other internal organs (not shown). Label-intensity registration (C) resulted in a more accurate overlay of the transformed subject image over the template image. In wildtype subjects, both registration approaches produce nearly identical results (B, D). The scaling associated with the initial linear transforms differ between registration approaches in both knockout and wildtype subjects (E-H), but the difference in scaling is much more apparent in knockouts (E, G). This representative wildtype subject did not contribute to the template image.

#### Label-informed image registration improves *Gli2^−/-^* registration

To qualitatively assess the transformations generated by intensity-only registration and label-informed intensity registration, each subject (yellow) was transformed into the coordinate system of the template (*i.e.,* template space) and overlaid onto the template (cyan) with the alpha (opacity) channel of both images set to 0.5 in 3D Slicer. As such, anatomical regions with a poor overlap of the transformed subject over the template appear as yellow or cyan, and areas of overlap appear green.

Intensity-only registration of most knockout did not result in accurate overlays over the template (Figure 3A). In contrast, label-informed intensity registration resulted in a much closer match between knockout subjects and the template (Figure 3C). In wildtype subjects, intensity-only and label-intensity registration produced similar overlays (Figure 3B,D).

The initial linear transforms calculated in the two registration approaches produce different scaling results in both knockout and wildtype subjects (Figure 3E-H). In intensity-only registration, knockout subjects were scaled to approximate the template image (Figure 3E,F) and then deformable transforms locally expanded and shrank the subject image (Figure 3G,H). In contrast, in the label-intensity registration, many of the knockout subjects fit around the template (Figure 3E,G) and the deformable transforms primarily shrank the image to align with the template. Consequently, for this dataset, the deformable transforms from label-informed intensity registration must be analyzed together with the linear transforms, and subsequent statistical analyses were conducted on the combined forward (subject to template space) transforms.

#### Tensor-based morphometry (TBM) yields greater statistical power

Tensor-based morphometry (TBM) compares the Jacobian determinants calculated from transformations to identify statistically significant voxel-wise differences in local volume (Ashburner & Friston 2000). Significantly different Jacobian values between knockout and wildtype subjects (FDR < 0.05, t-test) are shown as a heatmap overlaid over the template image (Figure 4). The magnitudes in the heatmap represent the magnitudes of the Jacobian determinant; negative Jacobian values indicate local volume reduction in knockout subjects relative to wildtypes and positive Jacobian values indicate local volume expansion in knockout subjects compared to wildtypes.

**Figure 4.**
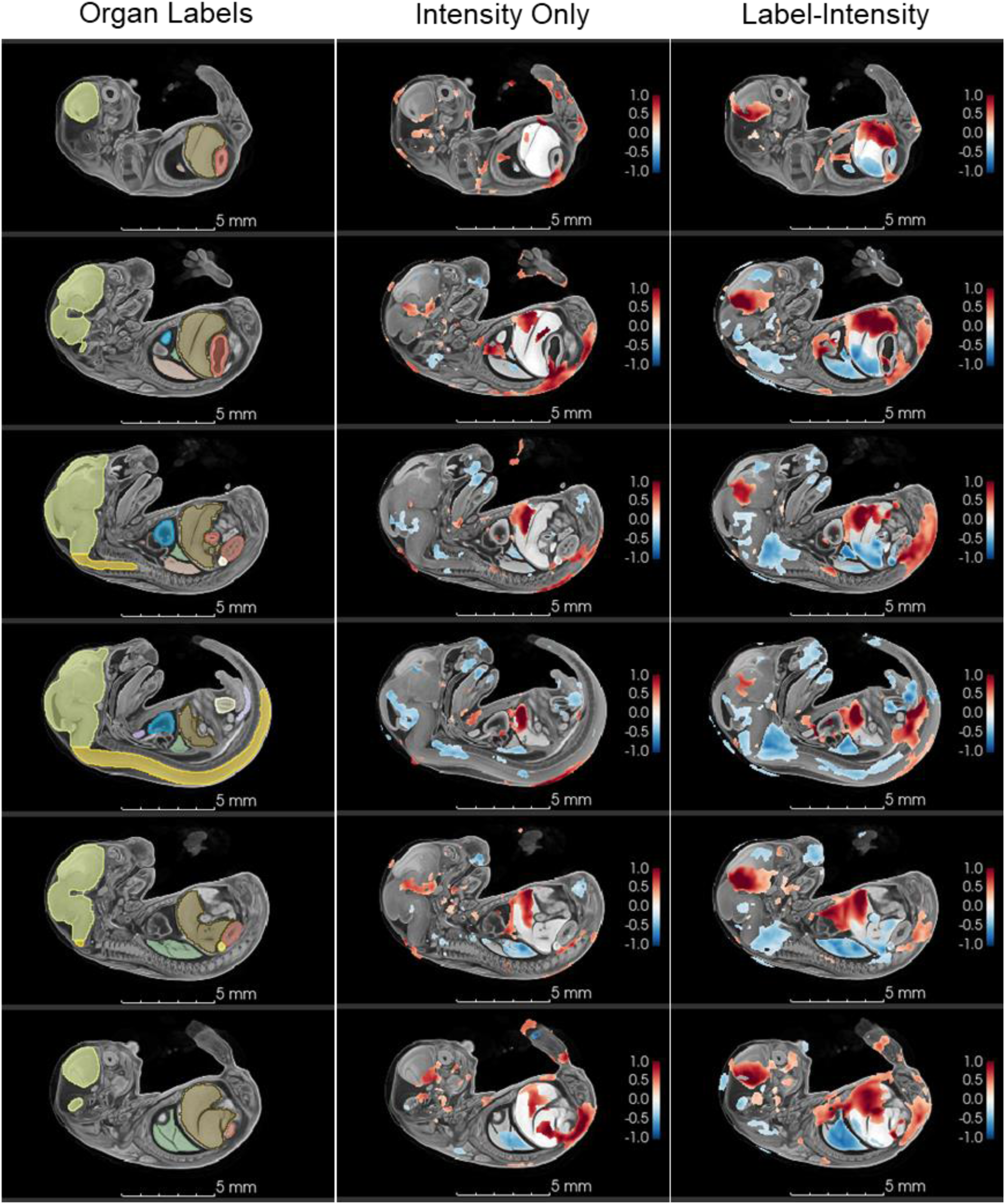
TBM heatmaps shown in serial sagittal slices in the template image. (left) Organ labels used in label-informed image registration; (middle) TBM result based on intensity-only registration; (right) TBM result based on label-informed image registration. Label-informed TBM showed a 50% increase in statistically significant (FDR < 0.05, t-test) Jacobian determinants compared to intensity-only TBM and a 45% increase in the range of Jacobian values. Label-informed TBM shows larger regions and magnitudes of volume reduction in the face, neck, lungs, liver, spine, bladder, and vesicorectal septum and volume expansion in the brain, thoracic cavity, liver, spine, and retroperitoneum. For comparative purposes, the heatmaps only show Jacobian values between -1 and 1.

TBM using label-informed image registration identified 50% more significantly different Jacobian values than TBM using intensity-only registration. Of the 1,255,976 voxels in the template image, tensor-based morphometry (TBM) using label-informed image registration identified 159,129 voxels as having significantly different Jacobian values while TBM using intensity-only registration only identified 103,707 (p < 0.05) (Figure 4).

The range of Jacobian magnitudes also increased in label-informed image registration compared to intensity-only registration. Jacobian values calculated from intensity-only registration transforms ranged from -0.99 to 1.94 whereas Jacobian values calculated from label-informed intensity registration transforms was -0.96 to 2.34, marking a 45% increase in range and greater sensitivity to local volume differences.

#### TBM results reflect observable anatomy and subtle size and shape differences

Qualitative assessment of individual knockout subject anatomy shows that local volume differences detected with TBM reflect observable morphological differences between knockout and wildtype subjects. TBM with both registration methods are grossly consistent with organ volume measurements calculated from the manually-segmented organ labels (the ground-truth) (Table 2). Volumes of organ labels that are significantly smaller (p < 0.05) in knockout subjects, such as the bladder and lungs, show regions with negative Jacobian values. Likewise, volumes of organ labels that are significantly larger (p > 0.05) in knockout subjects, such as the heart ventricles and liver, show positive Jacobian values (Table 2, Figure 4). However, intensity-only TBM failed to detect local volume differences in several regions that were detected in label-intensity TBM (Figure 4).

**Table 2.** Mean organ volumes for wildtype and knockout subjects calculated from manually-segmented organ labels.

| Organ | Wildtype | Knockout | T-test p-value |
| --- | --- | --- | --- |
| Adrenal (left) | 0.082 (0.009) | 0.053 (0.010) | < 0.001 |
| Adrenal (right) | 0.071 (0.011) | 0.050 (0.014) | 0.004 |
| Bladder | 0.445 (0.054) | 0.314 (0.105) | 0.006 |
| Brain | 28.826 (1.941) | 32.79 (5.313) | 0.062 |
| Heart | 1.601 (0.188) | 2.526 (1.111) | 0.038 |
| Kidney (left) | 0.554 (0.105) | 0.824 (0.326) | 0.041 |
| Kidney (right) | 0.627 (0.074) | 0.785 (0.274) | 0.129 |
| Liver | 17.603 (2.171) | 22.679 (4.530) | 0.011 |
| Lung (left) | 1.492 (0.253) | 0.990 (0.328) | 0.002 |
| Lung (right) | 3.118 (0.460) | 1.213 (0.338) | < 0.001 |
| Rectum | 0.136 (0.013) | 0.153 (0.065) | 0.459 |
| Spinal Cord | 7.287 (0.643) | 6.136 (1.090) | 0.017 |
| Stomach | 1.562 (0.171) | 1.838 (0.609) | 0.221 |
| Thymus (left) | 0.097 (0.015) | 0.107 (0.049) | 0.580 |
| Thymus (right) | 0.087 (0.017) | 0.101 (0.044) | 0.392 |

Within several organs that did not have significantly different label volumes (p > 0.05), label-informed TBM captured a combination of local volume increases and decreases, indicating differences in volumetric organ shape, which were not captured by intensity-only TBM (Figures 4,5). For example, multiple knockout subjects show anterodorsal displacement of the stomach with ventral displacement of the liver; in at least two subjects, this displacement is also associated with a diaphragmatic hernia (Figure 5). Volumes calculated from organ labels showed no significant difference in stomach volumes and a significant increase in liver volume in knockout subjects compared to wildtypes (Table 2). Intensity-only and label-informed TBM both showed that, compared to wildtypes, the knockout subjects had a volume expansion in a dorsal portion and a small volume reduction in an anterior portion of the stomach, and localized volume expansion in the ventral portion of the liver. However, label-intensity TBM also showed a volume reduction in the dorsal portion of the liver near the diaphragm, which was not captured in intensity-only TBM. The volume reduction captured by label-intensity reflects the ventral shift of the liver and anteroposterior compression of the dorsal liver, which is readily observable morphology in the knockout embryos.

**Figure 5.**
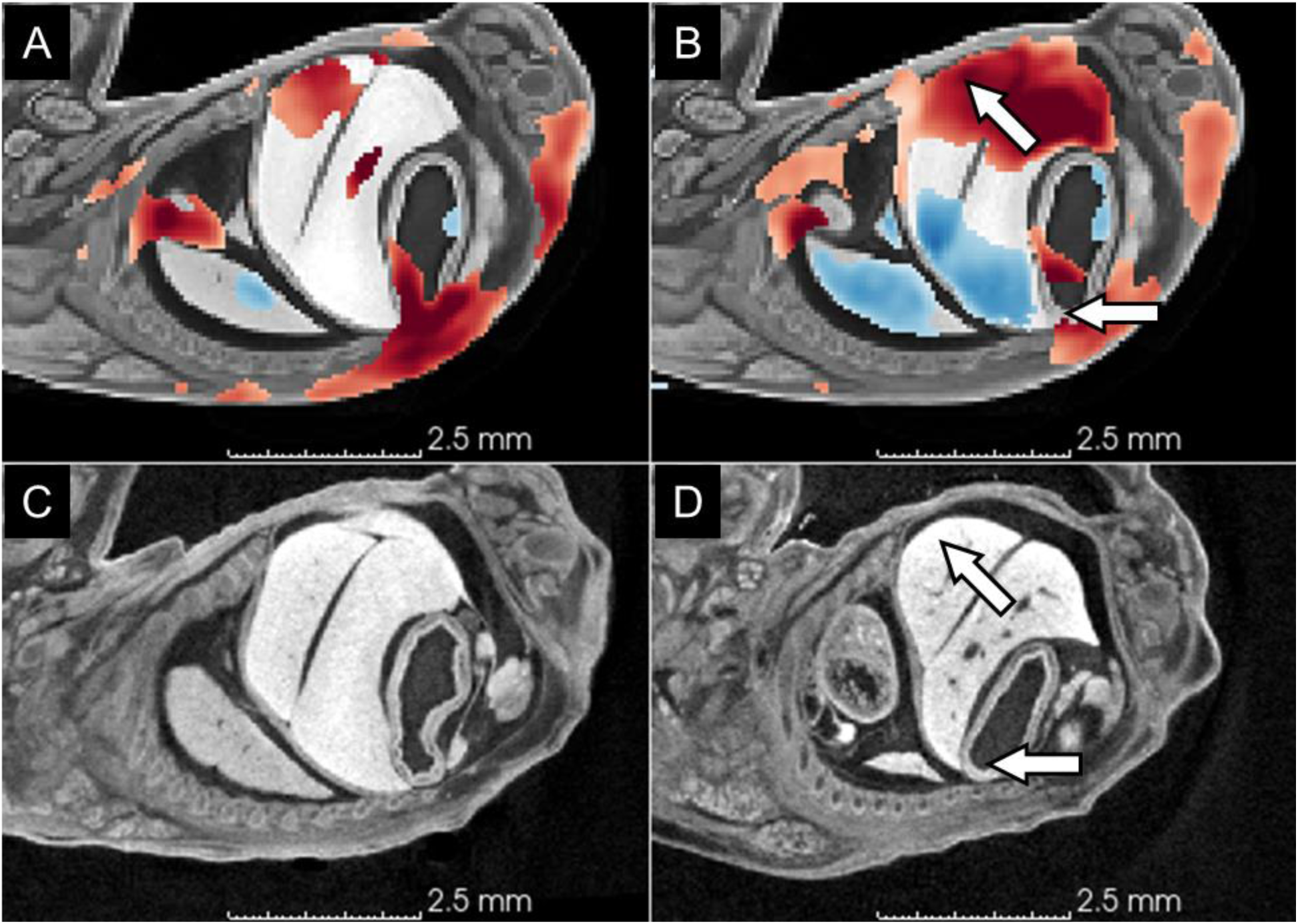
Anterodorsal displacement of the stomach. (A) Intensity-only TBM heatmap; (B) label-informed TBM heatmap; (C) a representative wildtype subject (Scan_0093) showing normal positions of the stomach and liver; (D) a representative knockout subject (Scan_0110) showing anterior displacement of the stomach and ventral displacement of the liver associated with longitudinal compression of the torso. White arrows indicate the anterior displacement of the stomach and ventral expansion of the liver in the knockout and the corresponding region in the TBM heatmap. Heatmaps use the same scale as in Figure 4. red = local volume expansion, positive Jacobians values, blue = local volume reduction in knockouts relative to wildtype subjects

A similar phenomenon is also observed in the spinal cord. The segment-based organ volumes show that knockout subjects have a smaller spinal cord than wildtypes (p < 0.05). However, the label volume alone gives no indication of the severe scoliosis observed in knockout subjects.

Intensity-only TBM shows small regions of ventral volume reduction in the cervical and lumbar regions of the spinal cord with a small region of volume expansion in the posterodorsal portion of the lumbar region closer to the rump. Label-intensity TBM heatmap shows much larger statistically significantly different regions as well as larger Jacobian magnitudes than intensity-only registration; and, the regions of volume difference more closely correspond with sharp bends in the spinal cord observed in knockout subjects (Figure 6).

**Figure 6.**
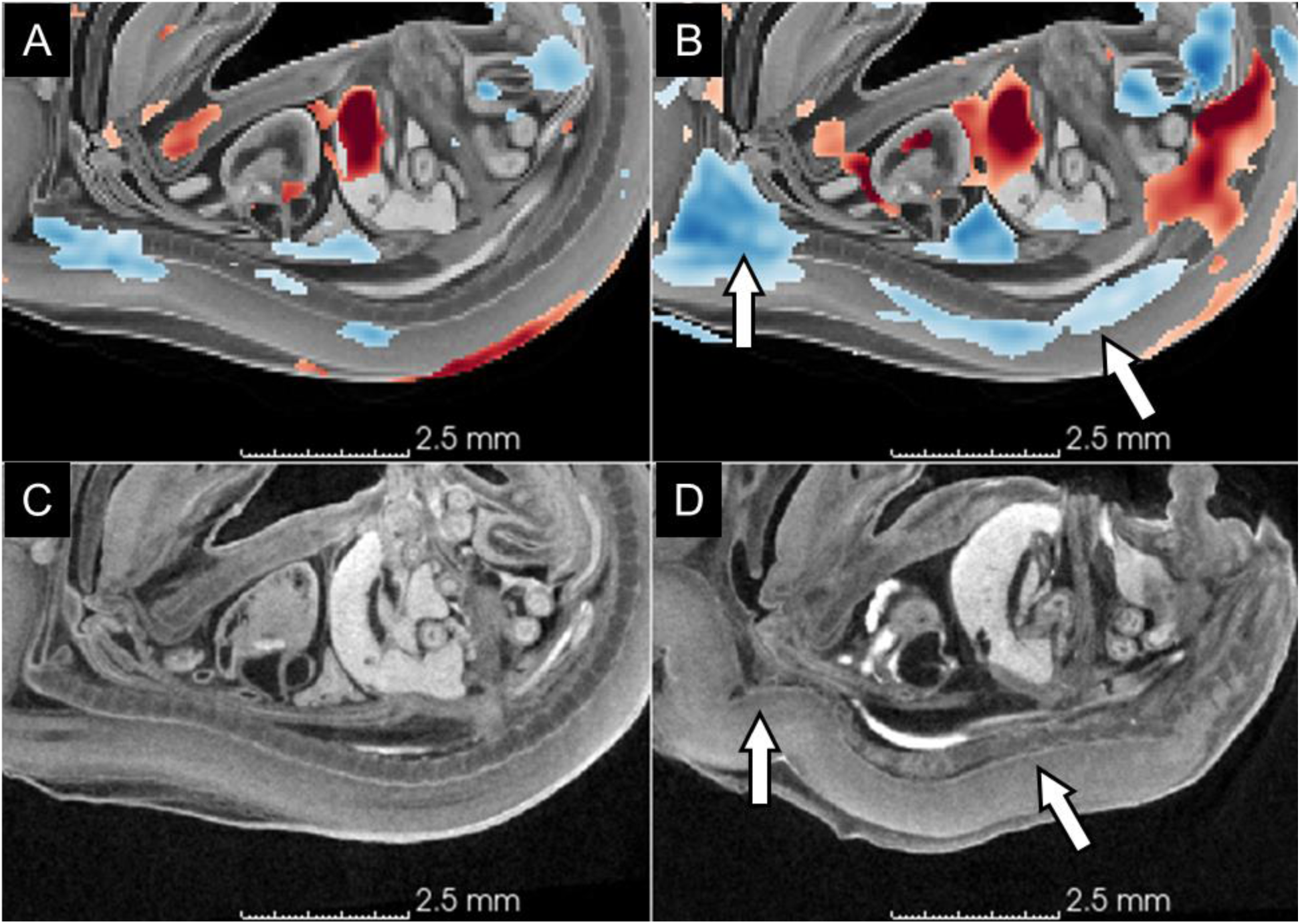
Scoliosis. (A) Intensity-only TBM heatmap; (B) label-informed TBM heatmap; (C) a representative wildtype subject (Scan_0093) showing smooth primary and secondary curvatures in the lumbar and thoracic regions; (D) a representative knockout subject (Scan_0105) showing severe bends and twists in the spinal cord and vertebral column. White arrows indicate the sharp bends in the knockout subject and corresponding negative (blue) Jacobian values in the TBM heatmap. Heatmaps use the same scale as in Figure 4. red = local volume expansion, positive Jacobians values, blue = local volume reduction in knockouts relative to wildtype subjects.

In body regions outside of the organ labels, label-intensity TBM also showed increased power and sensitivity compared to intensity-only registration. One example is a large region of volume expansion to the right of the heart ventricles (Figure 7). Intensity-only registration indicated only a small amount of volume expansion in this region, which includes both the right atrium of the heart and the right pleural space. Coronal slices of the thoracic cavity in knockout subjects clearly show that in the majority of subjects, the heart is displaced toward the left of the thoracic cavity, corresponding with the volume expansion observed in label-informed TBM (Figure 7).

**Figure 7.**
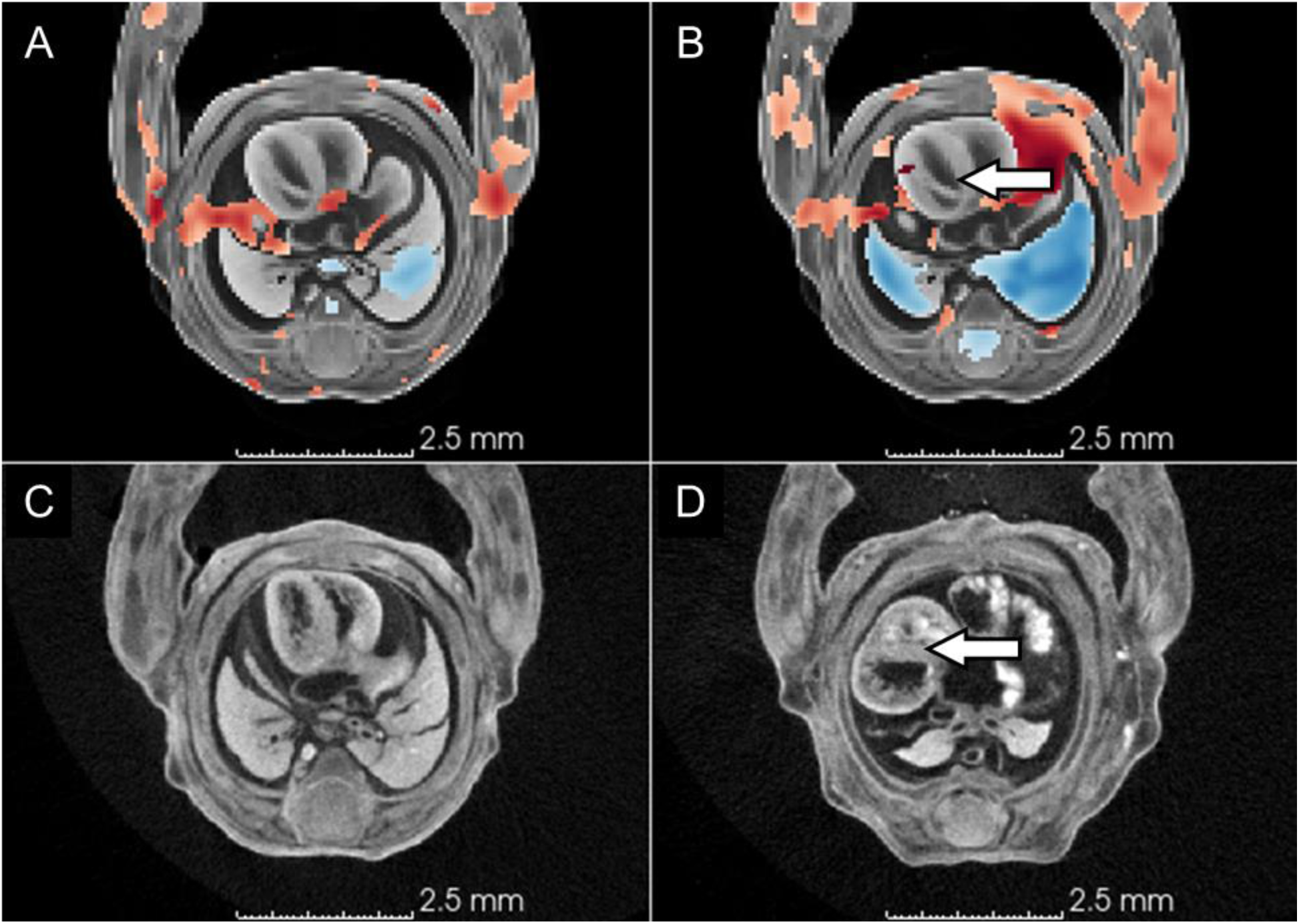
Displacement of the heart ventricles toward the left side of the thoracic cavity. (A) Intensity-only TBM heatmap; (B) label-informed TBM heatmap; (C) a representative wildtype subject (Scan_0093) showing the typical position of the heart ventricles within the thoracic cavity; (D) a representative knockout subject (Scan_0110) showing leftward displacement of the heart ventricles. White arrows indicate the leftward shift of the heart in the knockout subject and corresponding positive Jacobian values in the TBM heatmap. Heatmaps use the same scale as in Figure 4. red = local volume expansion, positive Jacobians values, blue = local volume reduction in knockouts relative to wildtype subjects.

Knockout subjects also show a qualitative reduction in the volume of both the left and right lungs, and only a single lobe in the right lung, which normally has four lobes (Figure 7C); this radical volume reduction is clearly shown in label-informed TBM, but intensity-only TBM only shows a small reduction in the centers of the right lung (Figure 7B) and left lung (Figure 4, not seen in the plane of Figure 7).

#### Label-informed PCA captures genotype-associated phenotypes

Principal component analysis (PCA) is a dimensionality reduction technique that allows visualization of morphospace and how individual subjects occupy morphospace. PCA on the combined inverse transforms provides a way to visualize how the template image is transformed to align with individual subject images.

PCA on the combined inverse transforms from intensity-only registration did not separate wildtype and knockout subjects on PC1 or PC2, or any other PC, even though there are obvious observable differences in knockout and wildtype subjects morphology due to severe scoliosis in the knockout subjects (Figures 2,8). However, PCA on the combined inverse transforms from label-informed intensity registration separated wildtype and knockout subjects on PC2 (Figure 8). Transformation of the template image along PC2 shows differences in the subjects’ spinal curvature (*i.e.,* scoliosis) between wildtype and knockout subjects (Figure 8). The PC2 transformations also show an increase in heart ventricle volume, differences in head shape and orientation consistent with observable subject phenotypes, and a reduction in the space between the bladder and rectum (associated with a vesicorectal fistula). PC1 primarily reflects overall subject size, as indicated by the PC1-associated transformations of the template image (Figure 8).

**Figure 8.**
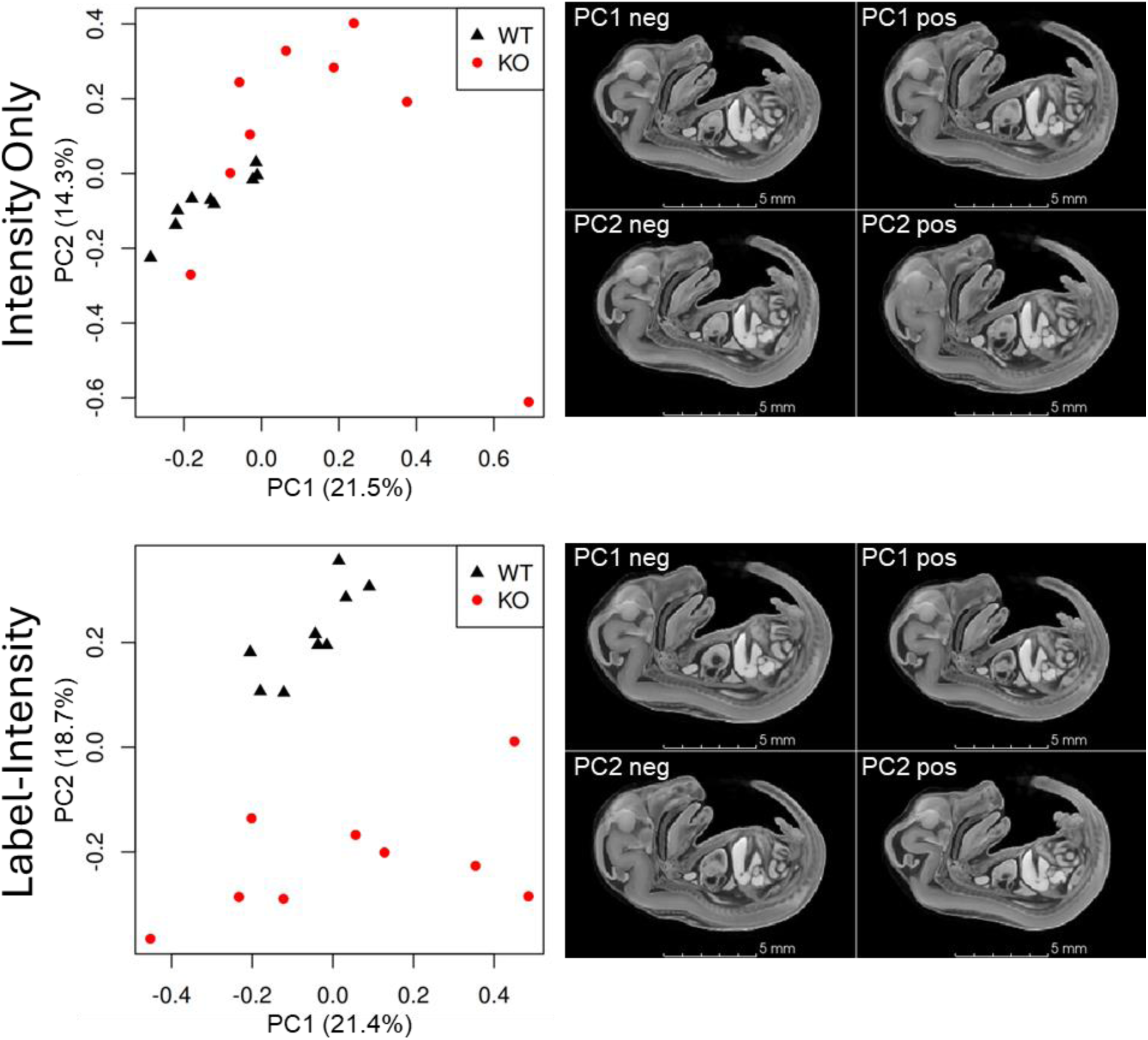
PC plots for combined inverse transforms from intensity-only registration (top) and label-intensity registration (bottom). Wildtype and knockout phenotypes separate along PC2 using transforms from label-informed registration, but do not separate on any PC in intensity-only registration. Transformation of the template image along PC2 shows overall morphology, including severe scoliosis, consistent with observable phenotypic differences (see Figure 2).

## Discussion

Label-informed image registration is a recently developed function in the ANTsX ecosystem that extends the capabilities of deformable image registration by incorporating anatomical knowledge via segmentation labels into the alignment process. Conceptually, this approach bridges the longstanding divide between atlas-based segmentation workflows and voxel-based morphometrics by allowing labeled structures to actively inform the registration, rather than serve merely as potential downstream outputs. Historically, intensity-based registration emerged as a powerful method for subtle morphometric analysis in neuroimaging, but difficulties can arise with large morphological variation, particularly in developmental and comparative studies where anatomy may be missing, shifted, or grossly deformed. Label-informed registration provides a solution by offering anatomically anchored initialization and refinement, thus enabling meaningful correspondence across highly divergent morphologies. This functionality is publicly available as part of the well-vetted, open-source ANTsR and ANTsPy libraries, which are widely used in biomedical imaging research and maintained with extensive community and peer-reviewed support. Importantly, the underlying nonlinear optimization is driven by the Symmetric Normalization (SyN) algorithm, the high-performing registration framework at the core of ANTsX, which has repeatedly demonstrated top-tier performance in comparative evaluations. By building on a foundation of robust, validated methods and extending them with anatomically-aware capabilities, this tool supports more accurate and interpretable morphometric analyses in cases where traditional registration techniques are prone to failure.

### Improved Image Registration and Morphometrics

Overt phenotypic differences in E15.5 *Gli2* knockout embryos have caused intensity-only registration to the wildtype template image to fail in large regions of the subject or for the entire image. *Gli2* knockout subjects differ in overall shape from wildtype subjects due to severe scoliosis and large displacements of internal organs (Figure 1). These overt differences have caused intensity-only registration of knockout subject images to the wildtype template image to fail in large regions of the subject or for the entire image. Label-informed image registration provided more accurate registration of these knockout subjects to a wildtype template image based on qualitative visual assessment of overlaid images. More accurate registration increased the power of downstream morphometric results using both tensor-based morphometry (TBM) and principal component analysis (PCA).

Because both wildtype and knockout subjects had better alignment with the template image, the calculated transformations (registration output) better reflect observable, qualitative morphological differences (Figures 4-7). TBM using both registration approaches showed local volume differences that were consistent with measured label volumes; but, TBM using label-informed intensity registration captured additional local volume differences not captured by intensity-only TBM that reflect observable differences in organ shape.

Likewise, PCA on intensity-only registration transforms did not show any genotype-associated phenotypic differences—wildtype and knockout subjects did not separate on any PC axis— despite readily apparent overt phenotypic differences. Because registration failed in several knockout subjects, the transforms used in this PCA do not reflect real morphological differences and the results are unreliable. In contrast, PCA on label-informed intensity registration transforms separated genotype-associated phenotypes on PC2, while PC1 primarily captured overall subject size. Thus, as expected, PCA on label-informed intensity registration transforms showed genotype is one of the main sources of morphological variation.

### Segmentation as a Registration Input

Manual image segmentation is a time-intensive process requiring substantial anatomical expertise (Rolfe et al. 2023). Just a few years ago, it would have been considered unfeasible to generate labels to use as an input for image registration for a larger number of images. Rather, labels have been one of the many potential derived data generated from finding correspondences in images through registration (*e.g.*, Xue et al. 2006). In atlas-based (registration-based) segmentation, a single template, or atlas, is expertly segmented, subjects are registered to the template image, and then the correspondences obtained through image registration are used to transform the atlas labels to match individual subjects (Horner et al. 2021; Wong et al. 2014; Xue et al. 2006).

Numerous semi-automated tools and deep learning models now exist to make segmentation relatively quick and easy (*e.g*., (Akinci D’Antonoli et al. 2025; He et al. 2024; Isensee et al. 2025; Rolfe et al. 2021; Rolfe et al. 2023; de Vente et al. 2025; Wasserthal et al. 2023). For example, the Segment Editor module in 3D Slicer and the SlicerMorph extension provide semi-automated tools to speed up image segmentation (Fedorov et al. 2012; Kikinis et al. 2014; Rolfe et al. 2021). Promptable deep learning-based frameworks like nnInteractive harness the benefits of deep learning while allowing users more control of segmentation (Isensee et al. 2025; de Vente et al. 2025). Several pre-trained models also provide fully-automated segmentation of multiple organs (Akinci D’Antonoli et al. 2025; Rolfe et al. 2023; Wasserthal et al. 2023). It is also now easier to train task-specific segmentation models from a small amount of training data through aggressive data augmentation (Tustison et al. 2024a,b). Because segmentations can now be generated relatively easily and independently of image registration, labels can now be used as an input for registration.

Many current anatomy-aware registration approaches are task-specific; that is, they perform well in the specific context for which they were designed for (*e.g.*, Azampour et al. 2024; Chilaparasetti et al. 2025; Hoffmann et al. 2024). The label-informed image registration function in the ANTsX ecosystem offers a general-purpose solution for anatomy-aware registration for a wide array of morphological variation. Because the labels are generated completely separately from the registration, they can be generated with any segmentation approach–manually, using semi-automated segmentation tools, or using task-specific deep learning models. This flexibility and customizability, allows users more options for defining regional correspondences to guide the registration.

#### Limitations of Label-Informed Intensity Registration

Label-informed intensity registration allows the registration of more diverse (or, geometrically dissimilar) morphologies and this greater diversity can add additional considerations when comparing individuals and populations. For example, there are many ways to choose a baseline for scaling subjects with different overall shapes depending on the morphological question (Dryden & Mardia 2016). Differences in overall shape between subjects or populations can also affect the scaling produced by linear transforms, and therefore the ability to use the linear transforms as a proxy for scale. In the present study, severe scoliosis causes *Gli2* knockout subjects to be relatively anteroposteriorly compressed compared to wildtype counterparts and to have a different angle between the head and torso (Figures 1,2). As a result, the linear transforms calculated using label-informed intensity registration increased the overall scale of knockout subjects to fit the subject around the template image, rather than approximate knockout subjects to the template (Figure 3). Then, the deformable transforms primarily shrank the image to fit the template. While this outcome is likely specific to the *Gli2* knockout phenotype and the specific labels selected for this study, it indicates that scaling from label-informed intensity registration may differ from other registration methods and between different morphologies, and that scaling may not be consistent between intensity image and label datasets.

While most registration methods are optimized to produce consistent results, label-informed image registration results can be biased by label choice, the distribution of labels across the image, and the relative weighting of each label. Selecting labels concentrated in only one region of an image could skew the registration toward that region and cause poor registration of other regions. Selected labels may also cause large or unrealistic transformations that are not relevant or informative. Additionally, labels must be present in all of the images being registered, which means that corresponding structures must be present in all the images. This limits, for example, the registration of subjects at later developmental stages to earlier developmental stages in which certain structures may not yet have developed.

### Potential Applications of Label-Informed Image Registration

#### Label-Guided Atlas-Based Segmentation

Depending on the research question, deep learning-based segmentation may not be an adequate replacement for atlas-based segmentation. While it has improved significantly, deep learning-based segmentation has several inherent limitations. Supervised training can require large amounts of well-annotated training data, which may be difficult to generate (Wang et al. 2021). Additionally, inference is often poor for novel data that was not included in the training dataset (Billot et al. 2023). And, unlike atlas-based segmentation which results in smooth transformations of atlas labels, deep learning-based segmentation can introduce small islands and other noise that can be difficult to detect. In many cases, it is likely that registration accuracy can be greatly improved with the introduction of only a small number of initial labels in a label-informed registration approach. Then, the more accurate transforms could be used to transfer additional labels from an atlas to subjects using an atlas-based segmentation approach.

#### Developmental and Comparative Morphometrics

Landmark-based geometric morphometric approaches have become commonplace in ontogenetic and inter-species comparative studies (*e.g.,* Churchill et al. 2018; Coombs et al. 2022; Devine et al. 2022; Klingenberg & Marugan-Lobon 2013; Young et al. 2014; Zelditch et al. 2012). But, landmarks often provide relatively sparse sampling of a volumetric (voxel) image and capture only a small portion of the overall shape variation (Deserno 2001; Roth 1993). Surface-based morphometrics can also provide detailed shape analysis, but do not capture the internal structure of anatomical features (*e.g.*, Rolfe & Maga 2023). Previously, registration-based morphometrics has been restricted to subjects with a high degree of morphological similarity and has been difficult to apply in comparative contexts. Label-informed image registration offers a way to conduct volumetric morphometric studies on a broader range of phenotypic diversity including different species and different developmental stages in which homologous regions can be identified.

## Contributions

R.A.R, N.J.T., and A.M.M conceptualized the study and data analysis, interpreted results, and revised and edited the manuscript. R.A.R. and N.J.T. drafted the initial manuscript; R.A.R. acquired the data, performed data analysis, and created the figures and tables; N.J.T. developed the labelImageRegistration() function; A.M.M. acquired the funding.

## Acknowledgements

We thank our colleagues at Seattle Children’s Research Institute and the University of Washington for their contributions to this project: D. Beier, S. Gombart, E. Vincent, and S. Houghtaling for mouse husbandry, genetics, and embryo collection; M. Bell and D. Mao for assistance with specimen preparation and micro-CT scanning; and S. Rolfe, C. Zhang, O. Thomas, and E. Nichols for valuable discussion and feedback on this project. 3D scanning of subjects was conducted at the SCRI MicroCT Imaging Facility (<u>RRID:SCR_024678</u>), using Bruker Skyscan 1272 microCT that was funded by an NIH shared instrument grant (<u>S10OD032302</u>). This work was supported by funding from the National Institutes of Health (<u>NICHD/P01HD104435</u> and <u>NIDCR/R03DE031313)</u>.

## Citations

Abouheif E. 1997. Developmental genetics and homology: a hierarchical approach. Trends in ecology & evolution 12:405–408.

Akinci D’Antonoli T, Berger LK, Indrakanti AK, Vishwanathan N, Weiss J, Jung M, Berkarda Z, Rau A, Reisert M, Küstner T, et al. 2025. TotalSegmentator MRI: Robust Sequence-independent Segmentation of Multiple Anatomic Structures in MRI. Radiology 314:.

Ashburner J, Friston KJ. 2000. Voxel-Based Morphometry—The Methods. NeuroImage 11:805–821.

Avants BB. 2024. ANTsR: ANTs in R: Quantification Tools for Biomedical Images.

Avants BB, Tustison N, Song G. 2009. Advanced normalization tools (ANTS). The Insight Journal 2:1–35.

Avants BB, Tustison NJ, Song G, Cook PA, Klein A, Gee JC. 2011. A reproducible evaluation of ANTs similarity metric performance in brain image registration. NeuroImage 54:2033–2044.

Azampour MF, Tirindelli M, Lameski J, Gafencu M, Tagliabue E, Fatemizadeh E, Hacihaliloglu I, Navab N. 2024. Anatomy-aware computed tomography-to-ultrasound spine registration. Medical Physics 51:2044–2056.

Bateson W. 1894. Materials for the Study of Variation: Treated with Especial Regard to Discontinuity in the Origin of Species. 1st ed.

Billot B, Greve DN, Puonti O, Thielscher A, Van Leemput K, Fischl B, Dalca AV, Iglesias JE. 2023. SynthSeg: Segmentation of brain MRI scans of any contrast and resolution without retraining. Medical Image Analysis 86:102789.

Boyer DM, Puente J, Gladman JT, Glynn C, Mukherjee S, Yapuncich GS, Daubechies I. 2015. A New Fully Automated Approach for Aligning and Comparing Shapes. The Anatomical Record 298:249–276.

Chen M, Tustison NJ, Jena R, Gee JC. 2023. Image registration: Fundamentals and recent advances based on deep learning. Machine Learning for Brain Disorders435–458.

Chilaparasetti AN, Thai A, Gao P, Xu X, Gopi M. 2025. RegBoost: Enhancing mouse brain image registration using geometric priors and Laplacian interpolation. NeuroImage 305:120981.

Churchill M, Geisler JH, Beatty BL, Goswami A. 2018. Evolution of cranial telescoping in echolocating whales (Cetacea: Odontoceti). Evolution 72:1092–1108.

Coombs EJ, Felice RN, Clavel J, Park T, Bennion RF, Churchill M, Geisler JH, Beatty B, Goswami A. 2022. The tempo of cetacean cranial evolution. Current Biology 32:2233–2247.

Cooper WJ, Albertson RC. 2008. Quantification and variation in experimental studies of morphogenesis. Developmental Biology 321:295–302.

Deserno TM. 2001. Biomedical Image Processing (Biological and Medical Physics, Biomedical Engineering).

Devine J, Vidal-García M, Liu W, Neves A, Lo Vercio LD, Green RM, Richbourg HA, Marchini M, Unger CM, Nickle AC, et al. 2022. MusMorph, a database of standardized mouse morphology data for morphometric meta-analyses. Scientific Data 9:230.

Dickinson ME, Flenniken AM, Ji X, Teboul L, Wong MD, White JK, Meehan TF, Weninger WJ, Westerberg H, Adissu H, et al. 2016. High-throughput discovery of novel developmental phenotypes. Nature 537:508–514.

Dryden IL, Mardia KV. 2016. Statistical shape analysis: with applications in R. John Wiley & Sons.

Fedorov A, Beichel R, Kalpathy-Cramer J, Finet J, Fillion-Robin J-C, Pujol S, Bauer C, Jennings D, Fennessy F, Sonka M, et al. 2012. 3D Slicer as an image computing platform for the Quantitative Imaging Network. Magnetic Resonance Imaging 30:1323–1341.

Goswami A, Clavel J. 2025. Morphological evolution in a time of phenomics. Paleobiology 51:195–213.

Groza T, Gomez FL, Mashhadi HH, Muñoz-Fuentes V, Gunes O, Wilson R, Cacheiro P, Frost A, Keskivali-Bond P, Vardal B, et al. 2023. The International Mouse Phenotyping Consortium: comprehensive knockout phenotyping underpinning the study of human disease. Nucleic Acids Research 51:D1038–D1045.

Hallgrímsson B, Jamniczky HA, Young NM, Rolian C, Schmidt-Ott U, Marcucio RS. 2012. The generation of variation and the developmental basis for evolutionary novelty. Journal of Experimental Zoology Part B: Molecular and Developmental Evolution 318:501–517.

He Y, Camaiti M, Roberts LE, Mulqueeney JM, Didziokas M, Goswami A. 2024. Introducing SPROUT (Semi-automated Parcellation of Region Outputs Using Thresholding): an adaptable computer vision tool to generate 3D segmentations.

Hoffmann M, Hoopes A, Greve DN, Fischl B, Dalca AV. 2024. Anatomy-aware and acquisition-agnostic joint registration with SynthMorph. Imaging Neuroscience 2:1–33.

Horner NR, Venkataraman S, Armit C, Casero R, Brown JM, Wong MD, Van Eede MC, Henkelman RM, Johnson S, Teboul L, et al. 2021. LAMA: automated image analysis for the developmental phenotyping of mouse embryos. Development 148:dev192955.

Isensee F, Rokuss M, Krämer L, Dinkelacker S, Ravindran A, Stritzke F, Hamm B, Wald T, Langenberg M, Ulrich C, et al. 2025. nnInteractive: Redefining 3D Promptable Segmentation [Internet].

Kandel BM, Cook PA, Tustison NJ, Muschelli J. 2024. ANTsRCore: Core Software Infrastructure for “ANTsR.”

Kikinis R, Pieper SD, Voxburgh K. 2014. 3D Slicer: a platform for subject-specific image analysis, visualization, and clinical support. In: Jolesz FA, editor. Intraoperative Imaging Image-Guided Therapy. New York, NY: Springer. p 277–289.

Klingenberg CP, Marugan-Lobon J. 2013. Evolutionary Covariation in Geometric Morphometric Data: Analyzing Integration, Modularity, and Allometry in a Phylogenetic Context. Systematic Biology 62:591–610.

Park H, Bai C, Platt K, Matise M, Beeghly A, Hui C c, Nakashima M, Joyner A. 2000. Mouse Gli1 mutants are viable but have defects in SHH signaling in combination with a Gli2 mutation. Development 127:1593–1605.

Patterson C. 1982. Morphological characters and homology. In: Joysey KA, Friday AE, editors. Problems of Phylogenetic Reconstruction. London: Academic Press. p 21–74.

Porto A, Rolfe S, Maga AM. 2021. ALPACA: A fast and accurate computer vision approach for automated landmarking of three-dimensional biological structures. Methods in Ecology and Evolution 12:2129–2144.

R Core Team. 2023. R: A Language and Environment for Statistical Computing. Vienna, Austria: R Foundation for Statistical Computing.

Rolfe S, Maga AM. 2023. DeCA: a dense correspondence analysis toolkit for shape analysis. International workshop on shape in medical imaging:259–270.

Rolfe S, Pieper S, Porto A, Diamond K, Winchester J, Shan S, Kirveslahti H, Boyer D, Summers A, Maga AM. 2021. SlicerMorph: An open and extensible platform to retrieve, visualize and analyse 3D morphology. Methods in Ecology and Evolution 12:1816–1825.

Rolfe SM, Whikehart SM, Maga AM. 2023. Deep learning enabled multi-organ segmentation of mouse embryos. Biology Open 12:bio059698.

Roth VL. 1984. On homology. Biological Journal of the Linnean Society 22:13–29.

Roth VL. 1991. Homology and hierarchies: problems solved and unresolved. Journal of Evolutionary Biology 4:167–194.

Roth VL. 1993. On three-dimensional morphometrics, and on the identification of landmark points. In: Marcus LF, Bello E, García-Valdecasas A, editors. Contributions to Morphometrics. Madrid: Monografías Museo Nacional de Ciencias Naturales CSIC. p 41–61.

Rutishauser R, Moline P. 2005. Evo-devo and the search for homology (“sameness”) in biological systems. Theory in Biosciences 124:213–241.

Scotland RW. 2010. Deep homology: A view from systematics. BioEssays 32:438–449.

Shubin N, Tabin C, Carroll S. 2009. Deep homology and the origins of evolutionary novelty. Nature 457:818–823.

Thompson D. 1917. On Growth And Form.

Tustison NJ, Chen M, Kronman FN, Duda JT, Gamlin C, Tustison MG, Kunst M, Dalley R, Sorensen SA, Wang Q, et al. 2024a. The ANTsX Ecosystem for Mapping the Mouse Brain.

Tustison NJ, Cook PA, Holbrook AJ, Johnson HJ, Muschelli J, Devenyi GA, Duda JT, Das SR, Cullen NC, Gillen DL, et al. 2021. The ANTsX ecosystem for quantitative biological and medical imaging. Scientific Reports 11:9068.

Tustison NJ, Yassa MA, Rizvi B, Cook PA, Holbrook AJ, Sathishkumar MT, Tustison MG, Gee JC, Stone JR, Avants BB. 2024b. ANTsX neuroimaging-derived structural phenotypes of UK Biobank. Scientific Reports 14:.

Van Valen LM. 1982. Homology and causes. Journal of Morphology 173:305–312.

de Vente C, Venkadesh KV, Ginneken B van, Sánchez CI. 2025. SlicerNNInteractive: A 3D Slicer extension for nnInteractive.

Wagner GP. 2014. Homology, Genes, and Evolution. Princeton: Princeton University Press.

Wang S, Li C, Wang R, Liu Z, Wang M, Tan H, Wu Y, Liu Xinfeng, Sun H, Yang R, et al. 2021. Annotation-efficient deep learning for automatic medical image segmentation. Nature Communications 12:.

Wasserthal J, Breit H-C, Meyer MT, Pradella M, Hinck D, Sauter AW, Heye T, Boll DT, Cyriac J, Yang S, et al. 2023. TotalSegmentator: Robust Segmentation of 104 Anatomic Structures in CT Images. Radiology: Artificial Intelligence 5:.

Wong MD, Dorr AE, Walls JR, Lerch JP, Henkelman RM. 2012. A novel 3D mouse embryo atlas based on micro-CT. Development 139:3248–3256.

Wong MD, Maezawa Y, Lerch JP, Henkelman RM. 2014. Automated pipeline for anatomical phenotyping of mouse embryos using micro-CT. Development 141:2533–2541.

Wong MD, Spring S, Henkelman RM. 2013. Structural Stabilization of Tissue for Embryo Phenotyping Using Micro-CT with Iodine Staining. PLoS ONE 8:e84321.

Xue Z, Shen D, Karacali B, Stern J, Rottenberg D, Davatzikos C. 2006. Simulating deformations of MR brain images for validation of atlas-based segmentation and registration algorithms. NeuroImage 33:855–866.

Young NM, Hu D, Lainoff AJ, Smith FJ, Diaz R, Tucker AS, Trainor PA, Schneider RA, Hallgrímsson B, Marcucio RS. 2014. Embryonic bauplans and the developmental origins of facial diversity and constraint. Development 141:1059–1063.

Zamyadi M, Baghdadi L, Lerch JP, Bhattacharya S, Schneider JE, Henkelman RM, Sled JG. 2010. Mouse embryonic phenotyping by morphometric analysis of MR images. Physiological Genomics 42A:89–95.

Zelditch M, Swiderski D, Sheets HD. 2012. Geometric morphometrics for biologists: a primer.

Zhang C, Porto A, Rolfe S, Kocatulum A, Maga AM. 2022. Automated landmarking via multiple templates. PLOS ONE 17:e0278035.

